# Short-chain fatty acids modulate the development and the cell surface molecule expression of DCs by epigenetic regulation

**DOI:** 10.64898/2025.12.23.696329

**Authors:** Weiting Zhao, Kazuki Nagata, Risako Akiyama, Yuki Yamazaki, Hiroto Kouda, Ryosuke Miura, Kenta Ishii, Ryusei Tokita, Naoto Ito, Norimasa Yamasaki, Osamu Kaminuma, Chiharu Nishiyama

**Affiliations:** Department of Biological Science and Technology, Faculty of Advanced Engineering, Tokyo University of Science, 6-3-1 Niijuku, Katsushika-ku, Tokyo 125-8585, Japan; Department of Disease Model, Research Institute for Radiation Biology and Medicine, Hiroshima University, 1-2-3 Kasumi, Minami-ku, Hiroshima 734-8553, Japan

**Keywords:** butyrate, dendritic cell, histone deacetylase, *Itga4*, LPAM-1, PU.1, short-chain fatty acid

## Abstract

**Background:** Short-chain fatty acids (SCFAs) are produced by the gut microbiota as secondary metabolites during fermentation process of dietary fibers. Although SCFAs are beneficial for immuno-related diseases because they regulate the gene expression and functions of myeloid cells, the effects of SCFAs on the development of DCs remain unclear.

**Methods:** We analyzed the effect of SCFAs on the expression levels of surface proteins and mRNAs, and histone modification in Flt3L-induced bone marrow-derived DCs.

**Results:** SCFAs, particularly butyrate, regulated the expression of surface molecules on mouse bone marrow-derived dendritic cells (DCs): increases in MHCII, CD86, CD11b, and LPAM-1 (α4β7) levels and the ratio of CD11c^+^/PDCA-1^-^/B220^-^ conventional DCs (cDCs) to CD11c^+^/PDCA-1^+^/B220^+^ plasmacytoid DCs (pDCs). Experiments using inhibitors of histone deacetylase (HDAC) and Gi proteins, and GPR109A deficient mice indicated that butyrate regulated DCs by suppression of HDACs and not through a stimulatory effect on G protein-coupled receptors. Butyrate and the HDAC inhibitor, trichostatin A (TSA), increased the cDC/pDC ratio, surface LPAM-1 and *Itga4* mRNA, while the mRNA level of *Itgb7* was not affected by butyrate and was reduced by TSA. ChIP assays showed that butyrate and TSA increased histone acetylation in the *Itga4* and *Spi1* genes. Furthermore, the butyrate treatment increased the levels of *Spi1* mRNA and PU.1 protein and decreased those of *Spib*/SpiB in DCs. In knockdown (KD) experiments using siRNAs, the gene expression of *Itga4* was decreased by KD of *Spi1* or *Irf8*, and cDC/pDC ratio decreased by *Spi1* KD.

**Conclusions:** Butyrate controls the gene expression and development of DCs through epigenetic regulation and DC-related transcription factors.

## Introduction

Short-chain fatty acids (SCFAs), which are produced by the gut microbiota as secondary metabolites during the fermentation process of dietary fibers, play an important role in modulating host immune responses. Although immune responses effectively defend against cancer and pathogen infections, the excessive and/or persistent activation of immune cells causes inflammatory diseases. Previous studies have revealed the various beneficial effects of SCFAs for inflammatory and allergic diseases through their regulation of gene expression and functions in immune cells.

SCFAs stimulate the Gi-type G protein coupled receptors (GPCRs), GPR41, GPR43, and GPR109A as ligands. Acetate has been shown to ameliorate inflammatory bowel disease (IBD), rheumatoid arthritis, and asthma in mice by suppressing chemotaxis, phagocytosis, and ROS production by neutrophils and eosinophils via GPR43 ^1^. Propionate and acetate were found to up-regulate the expression of Foxp3 and IL-10 in intestinal regulatory T cells (Tregs) and prevent mouse colitis in a GPR43-dependent manner ^2^. Propionate regulated the migration and functions of DC/macrophage progenitors via GPR41 in bone marrow, which suppressed Th2-dependent allergic asthma in mice ^3^. GPR109A is specific to butyrate among SCFAs and is also a receptor for vitamin B3 (nicotinic acid), whereas GPR41 and GPR43 recognize a wide range of SCFAs ^4^. GPR109A deficiency has been shown to aggravate inflammatory colitis and colon cancer due to a reduction in the butyrate-induced anti-inflammatory effects of DCs ^5^.

SCFAs also function as histone deacetylase inhibitors (HDACi). Butyrate was found to enhance the histone acetylation and subsequent transactivation of the *Foxp3* gene in CD4^+^ T cells ^6, 7^, which is involved in the Treg-mediated suppression of inflammatory colitis in mice ^6^. In humans as well, butyrate promoted the in vitro differentiation of naïve CD4^+^ non-Tregs towards iTregs by regulating histone acetylation of Treg-related genes, including the *Foxp3* gene ^8^. A recent study reported that butyrate suppressed the IgE-dependent activation of mast cells (MCs) by down-regulating the gene expression of signal transduction-related kinases with HDACi activity ^9^. In addition to HDACi activity, the butyrate-GPR109A axis was found to play an important role in suppressing the IgE-dependent activation of MCs *in vivo* and *in vitro* in our study ^10^.

GPR109A and HDACi are both involved in the effects of butyrate on DCs. Butyrate and nicotinic acid have been shown to induce the expression of IL-10 and retinaldehyde dehydrogenase 1 (RALDH1) in mouse DCs and macrophages via GPR109A, which enhances Treg development and Treg-mediated protection against inflammation in the colon ^5^. In the case of human DCs, GPR109A and HDACi were both required for the butyrate-induced expression of RALDH1, which is involved in the polarization of naïve CD4^+^ T cells towards IL-10-producing type 1 Tregs ^11^. However, the role of SCFAs in the differentiation of DCs and the expression of DC-associated transcription factors remains largely unknown.

In the present study, we showed that SCFAs, particularly butyrate, regulated the expression of cell surface molecules on mouse DCs. When bone marrow-derived DCs (BMDCs) were treated with butyrate, the surface levels of antigen presentation-related molecules increased, integrin LPAM-1 appeared on the cell surface, and the population of CD11c^+^/PDCA-1^+^/B220^+^ plasmacytoid DC (pDC) disappeared. We examined the molecular mechanisms by which butyrate modulated the expression of these molecules by using inhibitors, chromatin immunoprecipitation (ChIP) assays, small interfering RNAs (siRNAs), and *Hcar2* (encoding GPR109A) deficient, or Flag-tagged *Spib* knock-in mice. We revealed the roles of epigenetic regulation and the DC-related transcription factor PU.1 in butyrate-mediated gene expression in DCs. Specifically, butyrate treatment increased histone acetylation levels of the *Spi1* (encoding PU.1) and *Itga4* (encoding α4 subunit of LPAM-1) genes, promoting PU.1/IRF8-dependent transcription of the *Itga4* gene.

## Methods

### Mice care and ethics committee approval

Mice were maintained under specific pathogen-free-conditions and all experiments using mice were performed following the guidelines of the Institutional Review Board of Tokyo University of Science. The present study was approved by the Animal Care and Use Committees of Tokyo University of Science: K22005, K21004, K20005, K19006, K18006, K17009, K17012, K16007, and K16010.

### Production of *Hcar2*-and *Spib*-mutated mice

The improved genome editing via oviductal nucleic acid delivery (*i*-GONAD) was performed as described ^12, 13^. Briefly, C57BL/6N female mice randomly selected regardless of estrous cycle stage were administered 2 mg (0.08 ml) of progesterone (Mochida Pharmaceutical, Tokyo, Japan or FUJIFILM Wako Pure Chemical, Osaka, Japan) once daily at 17:00-19:00 on days 1 and 2. On day 4, 0.5 mg (0.1 ml) of anti-inhibin monoclonal antibody (AIMA; Biogate, Gifu, Japan) was injected simultaneously ^12^. Females were paired with males (1:1) overnight from 17:00-19:00 on day 6, and the presence of a vaginal plug was checked the following morning (day 7). Plug-positive females were used for *i*-GONAD experiments on day 7.

For the *i*-GONAD method, single-guide RNA (sgRNA) was designed using the Benchling CRISPOR tool (https://www.benchling.com/crispr) and produced using a GeneArt Precision gRNA Synthesis Kit (#A29377; Thermo Fisher Scientific). Two sgRNAs were designed to recognize the sequences of the 5’ untranslated region (CAGAGTGCCTCACCTAGCGA) and the downstream region (GCCAAGGGGAACGGCACCGG) of the *Hcar2*-coding region, and those of the start codon of the *Spib* gene (AGCCTCCAGAGCAAGCATGG and TGCAGCCTCCAGAGCAAGCA), each matching 20 bp of the wild-type genomes. A CRISPR gene-editing cocktail containing 1 μg/μl Cas9 protein (#1081059; Integrated DNA Technologies, Inc., Skokie, IL, USA) and 30 μM of each sgRNA was freshly prepared in Opti-MEM medium (#31985-062; Thermo Fisher Scientific Inc., Waltham, MA, USA) ^12^. For introducing 3 x Flag tag into the *Spib* gene, 30-60 μM of single-strand DNA (tatataggagtggtggccagagccagcctAGCCTGCTCTGAACCACCGACTACAAAGACCATGACGGTGATTATAAAGATCATGACATCGATTACAAGGATGACGATGACAAGCTTGCTCTGGAGGCTGCACAgtaagtgggtaacccccaatcctgttc; Integrated DNA Technologies, Inc., Skokie, IL, USA) was also included.

The ovaries and oviducts of anesthetized females (around 15:00 of day 7) were exposed, and approximately 1.5 μl of the cocktail was injected into the lumen of the oviduct upstream of the ampulla region using a fine glass pipette connected to a mouthpiece. After injection, the oviduct was covered with a piece of Kimwipe tissue wetted with PBS and then pinched by a forceps-type electrode (#CUY650P5; NEPA GENE, Chiba, Japan). Electroporation was performed using NEPA21 (NEPA GENE). The electroporation conditions consisted of three sequential poring pulses (40 V, 5 ms, 50 ms intervals, 10% decay [± pulse orientation]) followed by three transfer pulses (10 V, 50 ms, 50 ms intervals, 40% decay [± pulse orientation]). On day 19 of gestation, offspring were delivered either by cesarean section or natural birth. Genotyping was performed by agarose gel electrophoresis of PCR-amplified target regions, followed by direct sequencing to detect CRISPR-Cas9-induced mutations. The primers used for *Hcar2*-knockout (KO) mouse genotyping were as follows: CAGTCGGCTTGCCTAAACTTATCC / ACCGAACTGCAAAGGGAAGAACTC (**Supplementary Fig. S1A**). The PCR product sizes of 2,412 bp and 340 bp indicated wild-type and KO alleles, respectively. The primers used for 3 x Flag-knock-in (*Flag-Spib*) mouse genotyping were as follows: TGCCATGTGACTAGAGGGTTGTGC / CCTTATCACCTGCCTCTGGGTGC (**Supplementary Fig. S1B**). The PCR product sizes of 274 bp and 344 bp indicated wild-type and knock-in alleles, respectively. Founder mice carrying KO alleles were backcrossed to the C57BL/6N strain for at least two generations.

### Cells

BMDCs comprising cDCs and pDCs were generated from the bone marrow cells of 7-week-old male C57BL/6J mice (Japan SLC, Hamamatsu, Japan) by cultivation in a RPMI-1640 (#R8758, Sigma-Aldrich, St. Louis, MO, USA, or #C11875500CP, Gibco, Thermo Fisher Scientific, Waltham, MA, USA)-based medium supplemented with 10% FBS (#F7524-500ML, Sigma Aldrich, St. Louis, MO, USA, or #FB-1003/500, Biosera, Nuaillé, France), 100 U/mL penicillin (#01711331, Meiji, Tokyo, Japan), 100 U/mL streptomycin (#01711679, Meiji), 10 mM HEPES (pH7.4) (#17514-15, Nacalai Tesque, Kyoto, Japan), 1% MEM non-essential amino acids (#06344-56, Nacalai Tesque), 1 mM sodium pyruvate (#P2256, Sigma-Aldrich), 100 μM 2-ME (#135-07522, Fujifilm Wako), and 100 ng/mL recombinant mouse Flt3L (#575508, BioLegend, San Diego, CA, USA) for 8 days. Instead of Flt3L, 20 ng/mL recombinant mouse GM-CSF (#576308, BioLegend) was used to induce BMDCs comprising cDCs.

Final concentrations of 0.5 mM butyric acid (#B103500, Sigma-Aldrich), 10 nM trichostatin A (TSA, #T6933, Fujifilm Wako), 0.5 mM nicotinic acid (#72309, Sigma-Aldrich), and/or 0.1 μg/mL pertussis toxin (PTX, # 516561, Calbiochem, San Diego, CA, USA) were added to the culture medium of BMDCs on days 4 and 6, ensuring that these substances were present from day 4 to day 8.

### Flow cytometric analysis

Cells were preincubated with 1 μg/mL Fc block (clone 93, BioLegend or clone 2.4G2, Tonbo Biosciences, San Diego, CA, USA) on ice for 5 min and were stained with PE/Cy7-labeled anti-mouse CD11c (clone N418, BioLegend, 1:1000), PerCP-labeled anti mouse-I-A/I-E (clone M5/114.15.2, BioLegend, 1:2000), PE-labeled anti-mouse CD86 (clone GL-1, BioLegend, 1:250), APC/Cy7-labeled anti-human/mouse CD11b (clone M1/70, BioLegend, 1:2000), APC-labeled anti-mouse PDCA1/CD317/BST2 (clone 927, BioLegend, 1:300), FITC-labeled anti-mouse/human CD45R/B220 (clone RA3-6B2, BioLegend, 1:300), and PE-labeled anti-mouse LPAM-1 (Integrin α4β7) (clone DATK32, BioLegend, 1:300) with 1 μg/mL DAPI (#11034-56, Nacalai Tesque) on ice for 15 min. After washing, stained cells were applied to a FACSLyric Analyzer (BD Biosciences, San Jose, CA, USA) or MACS Quant Analyzer (Miltenyi Biotec, Bergisch Gladbach, North Rhine-Westphalia, Germany) to collect data. Data analyses were performed with FlowJo software (Tomy Digital Biology, Tokyo, Japan).

### Quantitative PCR (qPCR) to measure mRNA levels

The RelilaPrep RNA Cell Miniprep System (#Z6012, Promega, Madison, WI, USA) or RNAzol RT Reagent (#RN190, Cosmo Bio, Tokyo, Japan) was used to extract total RNA from DCs. After the synthesis of cDNA from total RNA using ReverTraAce qPCR RT Master Mix (#FSQ-201, TOYOBO, Osaka, Japan), qPCR was performed by the StepOne Real-Time PCR System (Applied Biosystems, Waltham, MA, USA) with THUNDERBIRD SYBR qPCR Mix (#QPS-201, TOYOBO). The primers used were as follows.

*Itga4* (encoding integrin α4)

Forward; 5’- GCTTGTGAACCCAACTTCAT -3’

Reverse; 5’- CATTTGGAGCCATGCTAATC -3’

*Itgb7* (encoding integrin β7)

Forward; 5’- CAAGTCACCATGTGAGCAG-3’

Reverse; 5’- GTCAAGGTCACATTCACGTC -3’

*Spi1* (encoding transcription factor PU.1)

Forward; 5’- ATGTTACAGGCGTGCAAAATGG -3’

Reverse; 5’- TGATCGCTATGGCTTTCTCCA -3’

*Spib* (encoding transcription factor SpiB)

Forward; 5’- AGAGGACTTCACCAGCCAGA -3’

Reverse; 5’- AACCCCAGCAAGAACTGGTA -3’

*Irf4* (encoding transcription factor IRF4)

Forward; 5’- CCCCATTGAGCCAAGCATAA-3’

Reverse; 5’- GCAGCCGGCAGTCTGAGA-3’

*Irf8* (encoding transcription factor IRF8)

Forward; 5’- CCGGATATGCCGCCTATG -3’

Reverse; 5’- GCTGATGACCATCTGGGAGAA -3’

*Gapdh* (encoding GAPDH)

Forward; 5’- ACGTGCCGCCTGGAGAA -3’

Reverse; 5’- GATGCCTGCTTCACCACCTT-3’

### Chromatin immunoprecipitation (ChIP) assays

ChIP assays were performed by using anti-acetyl Histone H3 (Lys27) mouse IgG1 Ab (clone MABI0309 (CMA309), Cosmo Bio) or control mouse IgG (#02-6100, Invitrogen, Thermo Fisher Scientific) as previously described ^14^. Briefly, 1-2 x 10^6^ cells suspended in 1 mL PBS were fixed in 1% formaldehyde at room temperature for 10 min. After the addition of glycine to stop the fixation reaction, cells were washed, solubilized with SDS buffer in the presence of protease inhibitors, and then sonicated. IP was performed with the above-mentioned Abs and protein G agarose/Salmon Sperm DNA (#16-201, Upstate, Sigma-Aldrich). The nucleotide sequences of the primers to quantify chromosomal DNA were as follows.

*Itga4* -234/-156

Forward; 5’-TTGTGTTCCCCCAAGGGTTA-3’

Reverse; 5’-GACTCACACCCTGGCAGACA-3’

*Itga4* -659/-590

Forward; 5’-TGCTAAGTGAGGCAAGTCACAGA-3’

Reverse; 5’-AGCCAGGCAATAGCCTAACACA-3’

*Itga4* -2805/-2024

Forward; 5’-CACCCTGGGCTGGTGTTC-3’

Reverse; 5’-TTGCTTCTGGTAGCTCACTTAACTG-3’

### Transfection of small interfering RNA (siRNA)

We introduced siRNA into GM-CSF-induced BMDCs and Flt3L-induced BMDCs by electroporation using Nucleofector 2b (Lonza, Basel, Switzerland) with the Amaxa Mouse Dendritic Cell Nucleofector Kit (Lonza) and using Neon transfection system (Invitrogen, Carlsbad, CA, USA), respectively, according to the manufacturers’ instructions. *Spi1* siRNA (#MSS247676, Invitrogen), *Irf8* siRNA (#MSS236848, Invitrogen), and their GC content-matched appropriate controls in the Stealth RNAi siRNA Negative Control Kit (#12935100, Invitrogen) were used.

### Western blotting analysis

Western blotting analysis was performed as previously described^15^ using anti-PU.1 Ab (clone 9G7, #2258, Cell Signaling Technology), anti-Flag Ab (clone M2, #F1804, Sigma-Aldrich), anti-GAPDH Ab (clone 14C10, #2118, Cell Signaling Technology), anti-rabbit IgG Ab (#7074, Cell Signaling Technology), and anti-mouse IgG Ab (#7076, Cell Signaling Technology).

### Statistical analysis

A two-tailed Student’s t-test or a one-sample t-test was used to compare two samples, and a one-way ANOVA followed by Tukey’s multiple comparison test was used for comparisons among more than three samples.

## Results

### SCFAs regulate cell surface molecule levels on DCs and the ratio of cDCs to pDCs

To investigate the effects of SCFAs on the gene expression and development of DCs, we analyzed the cell surface levels of DC-related molecules on Flt3L-generated BMDCs. A DAPI staining analysis showed that the incubation with 0.5 mM SCFAs did not reduce the viability of BMDCs; this result was confirmed in multiple independent experiments (**Fig. 1A** top, and **Supplementary Fig. S2**). Treatments with butyrate increased the cell surface levels of MHCII, CD86, and CD11b, suggesting an activation state of DCs (**Fig. 1A** and **B**). In addition, the cell surface level of CD11c and the frequency of CD11c^+^ cells were reduced by the butyrate treatment (**Fig. 1A** and **C**). In addition to the decrease in surface level of CD11c following the SCFA treatment potentially reflecting the activation state of BMDCs ^16^, butyrate may have modulated the development of DCs because CD11c is a specific marker of DCs. Therefore, we investigated the frequencies of cDCs and pDCs in BMDCs using Abs against PDCA1 and B220 (**Fig. 1D**), and found that the butyrate treatment increased the frequency of cDCs and decreased that of pDCs (**Fig. 1E**). In BMDCs treated with various SCFAs (0.5 mM), an increase in cDCs and a decrease in pDCs were observed in the following order: butyrate > propionate > valerate, isobutyrate, isovalerate >> acetate ≒ 0 (**Fig. 1F**, and **Supplementary Fig. S3**).

**FIGURE 1.**
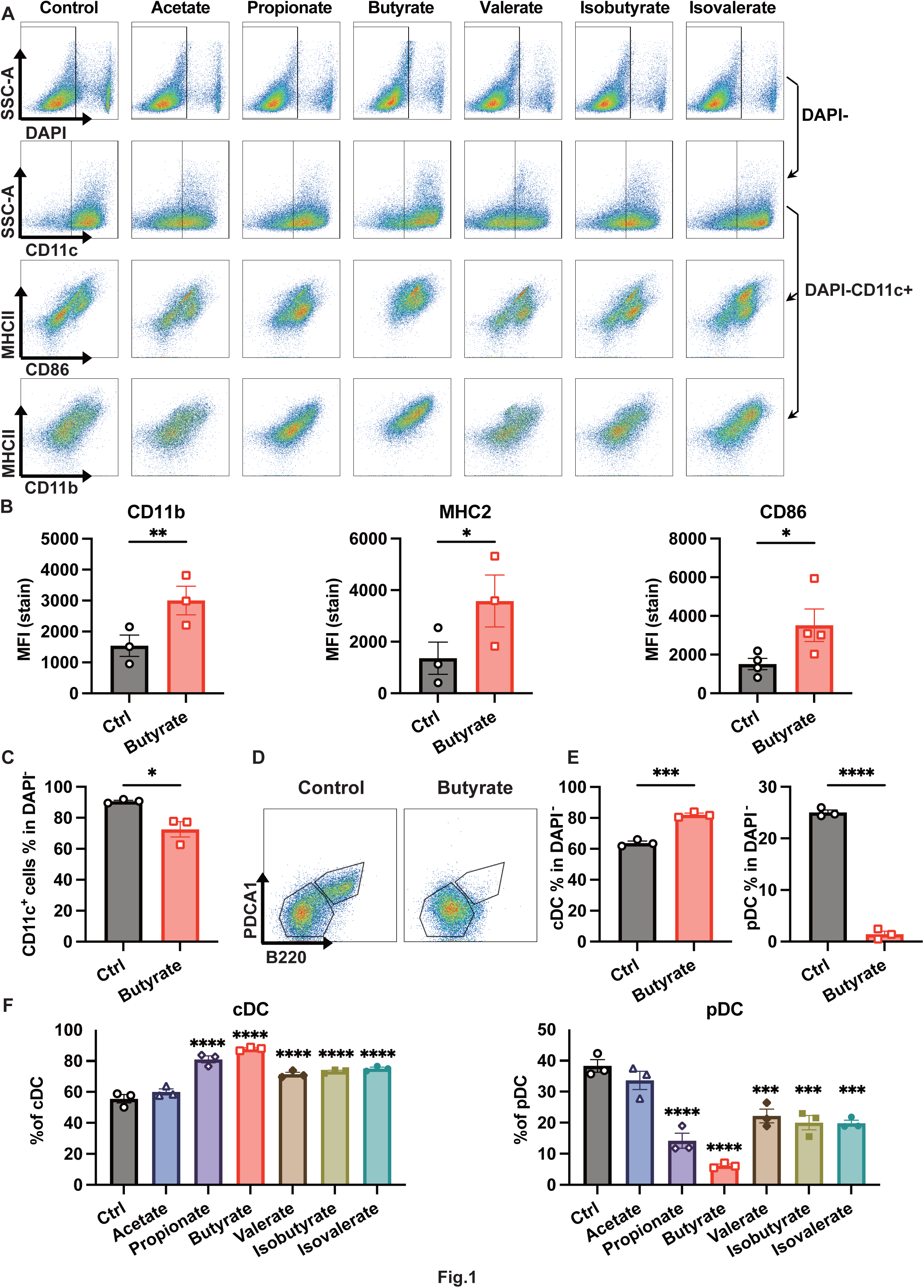
Effects of short-chain fatty acids (SCFAs) on expression levels of cell surface molecules in bone marrow-derived DCs (BMDCs). **A.** Dot plot profiles of Flt3L-induced BMDCs assessed by flow cytometry. The indicated SCFAs at a final concentration of 0.5 mM were added to the culture medium of BMDCs from day 4 to day 8 of the culture (**A**, **F**). A flow cytometric analysis was performed on day 8 (**A**-**F**). **B.** Mean fluorescence intensity (MFI) of BMDCs stained with APC/Cy7-labeled anti-CD11b, PerCP-labeled anti-I-A/I-E, or PE-labeled anti-CD86. BMDCs were generated in the presence or absence of 0.5 mM butyrate from days 4 to 8 (**B**-**E**). **C.** Percentage of CD11c^+^ cells in DAPI^-^ BMDCs. **D.** Typical dot plot profiles of the DAPI^-^/CD11c^+^ fraction of BMDCs stained with FITC-labeled anti-B220 and PE-labeled anti-PDCA1. **E.** Percentages of CD11c^+^/B220^-^/PDCA1^-^ conventional DCs (cDCs) (left) and of CD11c^+^/B220^+^/PDCA1^+^ plasmacytoid DCs (pDCs) (right) in the DAPI^-^ fraction. **F.** Percentages of cDCs (left) and pDCs (right) in BMDCs treated with SCFAs. Data represent the mean ± SEM of three independent experiments performed using biologically independent samples (**B**, **C**, **E**, **F**). The two-tailed paired Student’s *t*-test (**B**, **C**, **E**) and Tukey’s multiple comparison test (**F**) were used for statistical analyses. *, *p* < 0.05; **, *p* < 0.01; ***, *p* < 0.005.

We also found that the butyrate treatment significantly increased the cell surface level of LPAM-1 (α4/β7), an integrin that is involved in migration and localization in the intestinal mucosa, on BMDCs (**Fig. 2A**).

**FIGURE 2.**
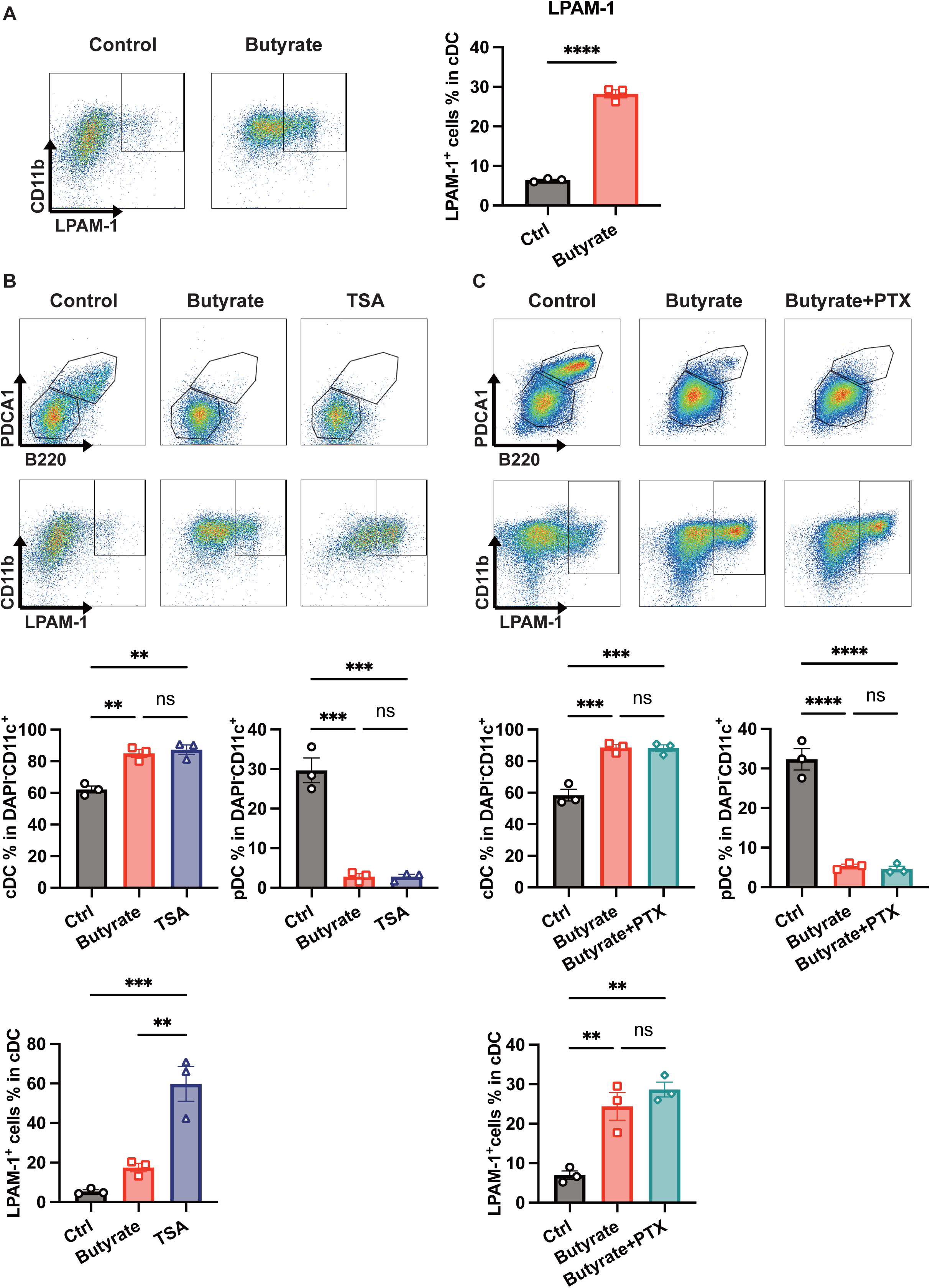
Effects of butyrate, trichostatin A (TSA), and pertussis toxin (PTX) on the conventional DC (cDC)/plasmacytoid DC (pDC) ratio and cell surface levels of integrin LPAM-1. **A.** Typical dot plot profiles of CD11b/LPAM-1 staining (left) and percentage of LPAM-1^+^ cells in cDC (right). **B.** Effects of butyrate and TSA on the percentages of cDCs, pDCs, and LPAM-1^+^ cells. **C.** Effects of PTX on butyrate-induced increases in cDCs and LPAM-1 levels and decreases in pDCs. Data represent the mean ± SEM of three independent experiments performed using biologically independent samples (right in **A**-**C**). The two-tailed paired Student’s *t*-test (**A**) and Tukey’s multiple comparison test (**B**, **C**) were used for statistical analyses. **, *p* < 0.01; ***, *p* < 0.005; ****, *p* < 0.001; ns, not significant.

Based on these results, we concluded that SCFAs regulate the expression of cell surface molecules on DCs, i.e. increases in MHCII, CD86, CD11b, and LPAM-1, and decreases in CD11c, PDCA1, and B220, indicating the stimulation of cDCs and modulation of the cDC/pDC ratio.

### Roles of HDACi activity and GPCR in effects of butyrate on DCs

SCFAs exhibit HDACi activities that regulate gene expression by increasing histone acetylation. The order of the effects of SCFAs on the cDC/pDC ratio (butyrate > propionate >> acetate ≒ 0) was consistent with the order of HDACi activities in SCFA-treated CD4^+^ T cells ^17^, and with the order of histone H3 acetylation levels in SCFA-treated BMDCs ^7^. To investigate the involvement of HDACi activity in the effects of SCFAs on DCs, we treated BMDCs with the HDACi TSA. TSA treatment significantly increased cDCs and decreased pDCs, as similar to the butyrate treatment (**Fig. 2B**). TSA also significantly increased the cell surface expression level of LPAM-1 on DCs more effectively than the butyrate treatment (**Fig. 2B**).

In addition to HDACi activity, Gi-type GPCRs, including GPR41, 43, and 109A, are the other major pathways of SCFAs, and DCs express GPR109A ^18^. We treated BMDCs with pertussis toxin (PTX), an inactivator of the Gi/o protein, to evaluate the involvement of GPCRs in the effects of SCFAs on DCs, and found that butyrate-induced increases in the cDC/pDC ratio and the cell surface expression of LPAM-1 were not affected by the PTX treatment (**Fig. 2C**). These results indicate that the effects of butyrate on the cell surface expression of PDCA1, B220, and LPAM-1 were mediated via HDACi activity, but not GPCRs.

### Effects of GPR109A deficiency and of GPR109A stimulation on DCs

In addition to the treatment with PTX, we conducted further experiments to investigate the involvement of GPR109A in the regulation of DCs, because acetate, a ligand for GPR41 and GPR43, did not cause apparent effects (**Fig. 1**), butyrate preferentially stimulates GPR109A among SCFA receptors ^19^, and GPR109A is highly expressed in murine DCs ^18, 20^ and human DCs (**Supplementary Fig. S4**). Then, we compared the effects of butyrate among BMDCs generated from the *Hcar2* (encoding GPR109A) KO mice and its control DCs, and found that GPR109A deficiency did not reduce the butyrate-induced increase in cDC/pDC ratio (**Fig. 3A**). Furthermore, we treated BMDCs with nicotinic acid, a ligand for GPR109A, which have been used to stimulate GPR109A on DCs ^5^, keratinocytes ^21^, Langerhans cells ^22^, and MCs ^10^, and found that the treatment with nicotinic acid did not affect cDC/pDC ratio (**Fig. 3B**).

**FIGURE 3.**
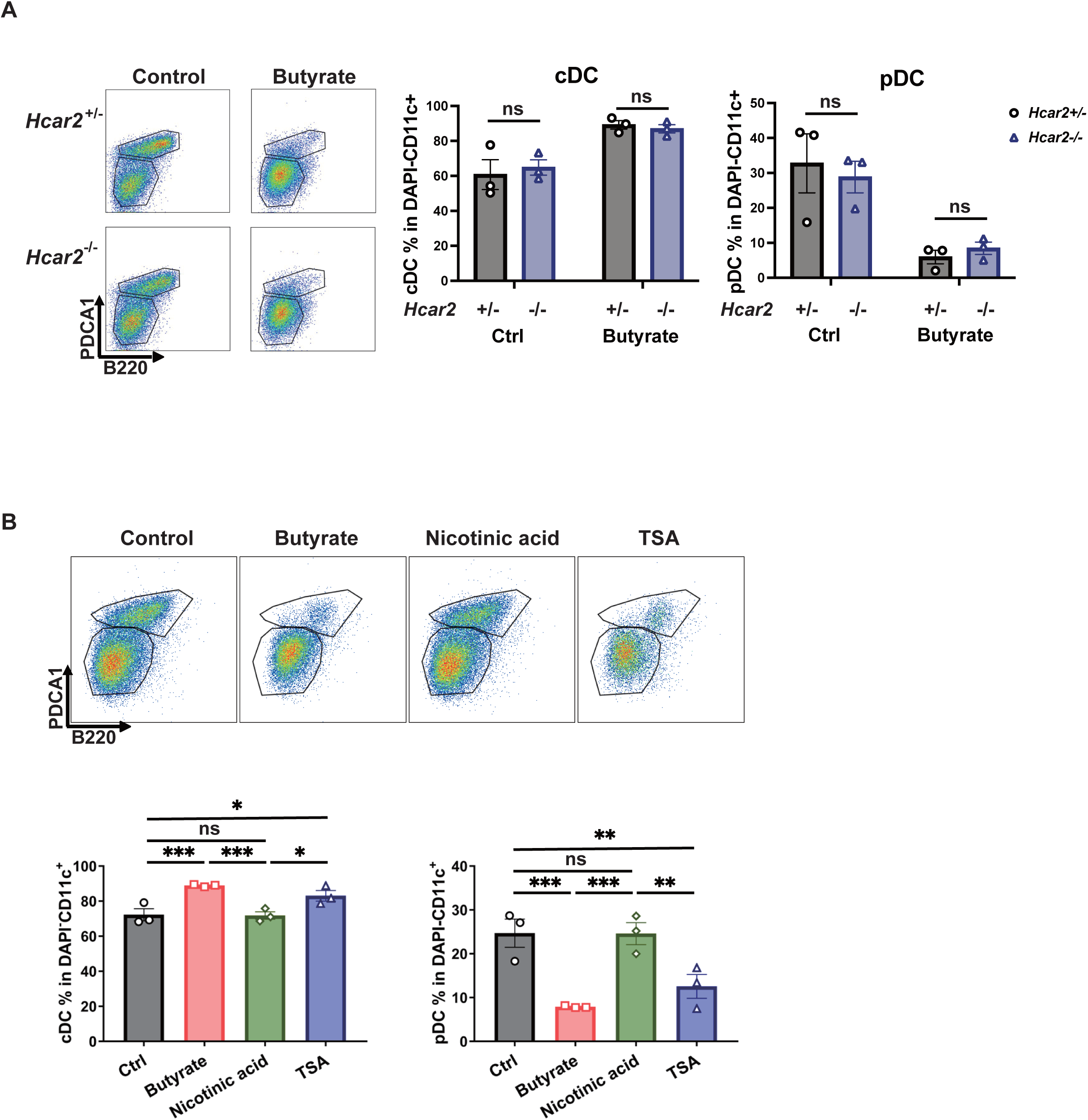
Effects of GPR109A deficiency and of GPR109A stimulation on DCs. **A.** Typical dot plot profiles stained with anti-B220 and anti-PDCA1 Abs in DAPI^-^/CD11c^+^ fractions of BMDCs generated from *Hcar2*^-/-^ and its control mice (**left**). Data represent the mean ± SEM of three independent experiments performed using biologically independent samples (**right**). **B.** Effects of GPR109A ligands and HDACi on the expression levels of cell surface molecules in DCs. Butyrate at a final concentration of 0.5 mM (**A**, **B**), 0.5 mM nicotinic acid and 10 nM TSA (**B**) were added to the culture medium of BMDCs from days 4 to day 8 of the culture. Data represent the mean ± SEM of three independent experiments performed using biologically independent samples (**B**). The two-tailed paired Student’s *t*-test (**A**) and Tukey’s multiple comparison test (**B**) were used for statistical analyses. *, *p* < 0.05; **, *p* < 0.01; ***, *p* < 0.005; ns, not significant.

### Butyrate and TSA increased histone acetylation levels on the Itga4 gene and Itga4 mRNA in DCs

To reveal the molecular mechanisms by which butyrate and TSA increased the cell surface level of LPAM-1, we measured the mRNA levels of *Itga4* and *Itgb7* in BMDCs. As shown in **Fig. 4A**, the mRNA level of *Itga4* significantly increased in DCs treated with butyrate or TSA, whereas *Itgb7* mRNA levels were unaffected by butyrate and were reduced by TSA, suggesting that the increased expression of cell surface LPAM-1 induced by butyrate or TSA reflected an increase in *Itga4* transcripts.

**FIGURE 4.**
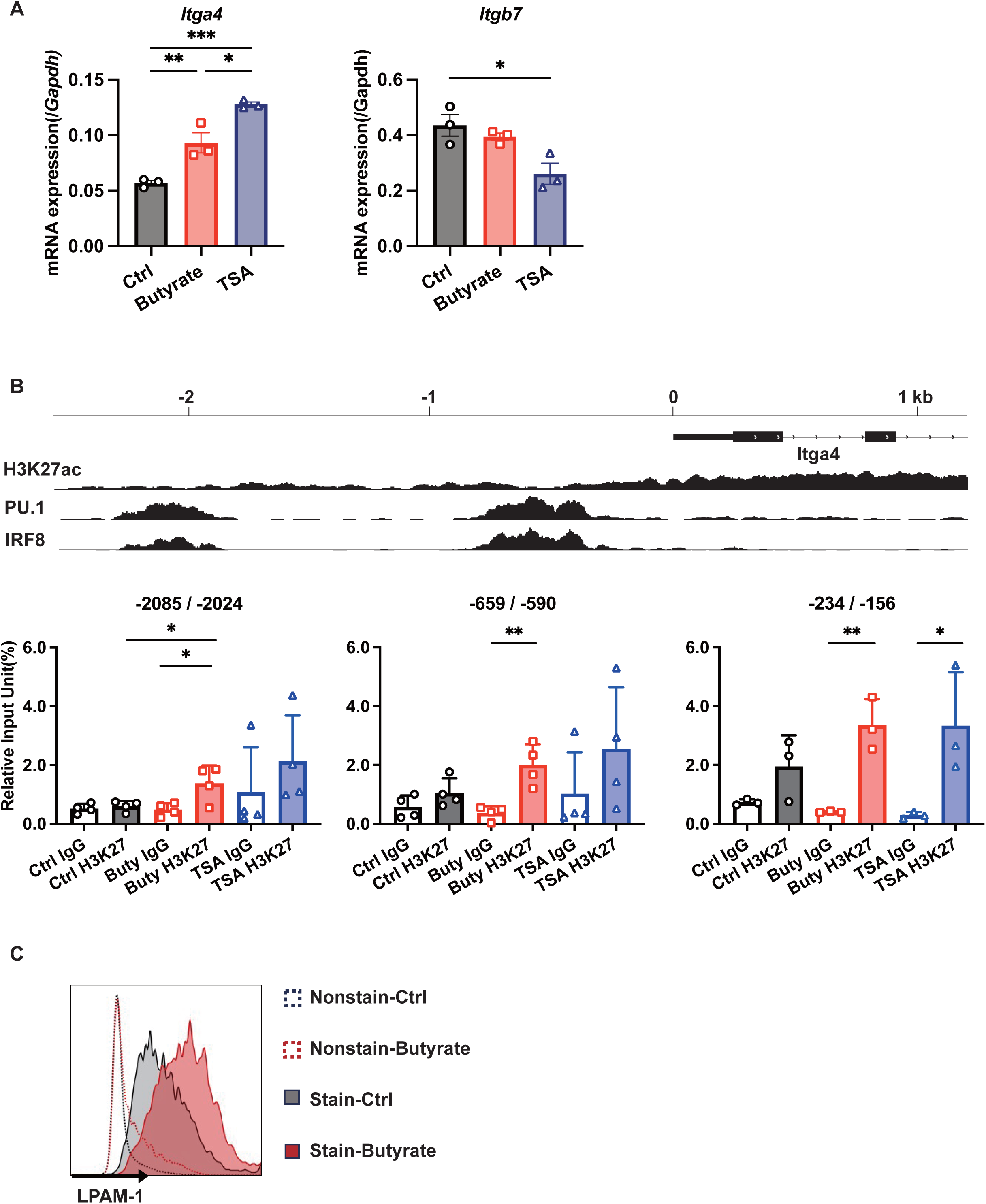
Butyrate and trichostatin A (TSA) up-regulated LPAM-1 expression on DCs by increasing *Itga4* mRNA expression levels via enhancements in the H3K27 acetylation of the *Itga4* gene. **A.** Messenger RNA levels of *Itga4* (encoding α4 subunit of LPAM-1) and *Itgb7* (β7 subunit of LPAM-1) in Flt3L-induced BMDCs. This figure shows the levels of *Itga4* mRNA and *Itgb7* mRNA normalized using *Gapdh* (a housekeeping gene) as a reference. **B.** H3K27 acetylation analyzed by a ChIP assay performed with an anti-acetyl Histone H3 (Lys27) Ab and its control IgG. Top: The profiles of H3K27 acetylation (accession number SRX1255004), PU.1-binding (SRX4909225), and IRF8-binding (SRX14466997) around the *Itga4* gene in cDCs were obtained via a ChIP-Atlas analysis (http://chip-atlas.org). Bottom: Relative input unit (%) of immunoprecipitated DNA assessed by qPCR targeting the indicated location on the *Itga4* gene. **C.** Butyrate increased the cell surface levels of LPAM-1 on DCs isolated from the mouse spleen. Data represent the mean ± SEM of three independent experiments performed using biologically independent samples (**A**, **B**). Tukey’s multiple comparison test (**A**, **B**) was used for statistical analyses. *, *p* < 0.05; **, *p* < 0.01; ***, *p* < 0.005.

To further investigate the effects of butyrate and TSA on histone acetylation around the *Itga4* gene, we performed ChIP assays. ChIP assays using an anti-acetyl H3K27 Ab and its control Ab revealed significant histone acetylation in the 5’-flanking region of the *Itga4* gene in butyrate-treated DCs, but not in untreated DCs (**Fig. 4B**). The TSA treatment also generally increased histone acetylation, although no significant difference was detected in two of three regions examined due to differences in the magnitude of changes in each independent experiment.

We also examined the effects of butyrate on LPAM-1 expression in cDCs by incubating whole splenocytes isolated from mice in the presence of butyrate. As shown in **Fig. 4C**, butyrate-treated cDCs expressed higher levels of LPAM-1 on their surface than non-treated cDCs.

### Effects of butyrate on the gene expression of DC-related transcription factors

*In silico* analyses of ChIP-seq data showed the apparent binding of the transcription factors PU.1 and IRF8 around butyrate-induced histone-acetylated regions on the *Itga4* gene in DCs (**Fig. 4B**). We then performed knockdown (KD) experiments using siRNA to investigate the involvement of PU.1 and IRF8 in *Itga4* gene expression in DCs. In this experiment, we aimed to evaluate the mRNA expression levels of transcription factors under conditions unaffected by fluctuations in the cDC/pDC ratio. Since LPAM-1 was detected in cDC population of Flt3L-induced BMDCs, we used GM-CSF-induced BMDCs, in which cDC were mainly generated. When *Spi1* (encoding PU.1) mRNA levels were significantly reduced by the transfection of *Spi1* siRNA, *Itga4* mRNA levels also decreased (**Fig. 5A**). In addition, *Irf8* KD significantly reduced *Itga4* mRNA levels (**Fig. 5B**). *Spi1* KD reduced *Spib* and *Irf8* mRNA levels (**Fig. 5A**), suggesting the role of PU.1 as a master transcription factor regulating other DC-related transcription factors, whereas *Irf8* KD did not affect *Spi1* and *Spib* mRNA levels (**Fig. 5B**).

**FIGURE 5.**
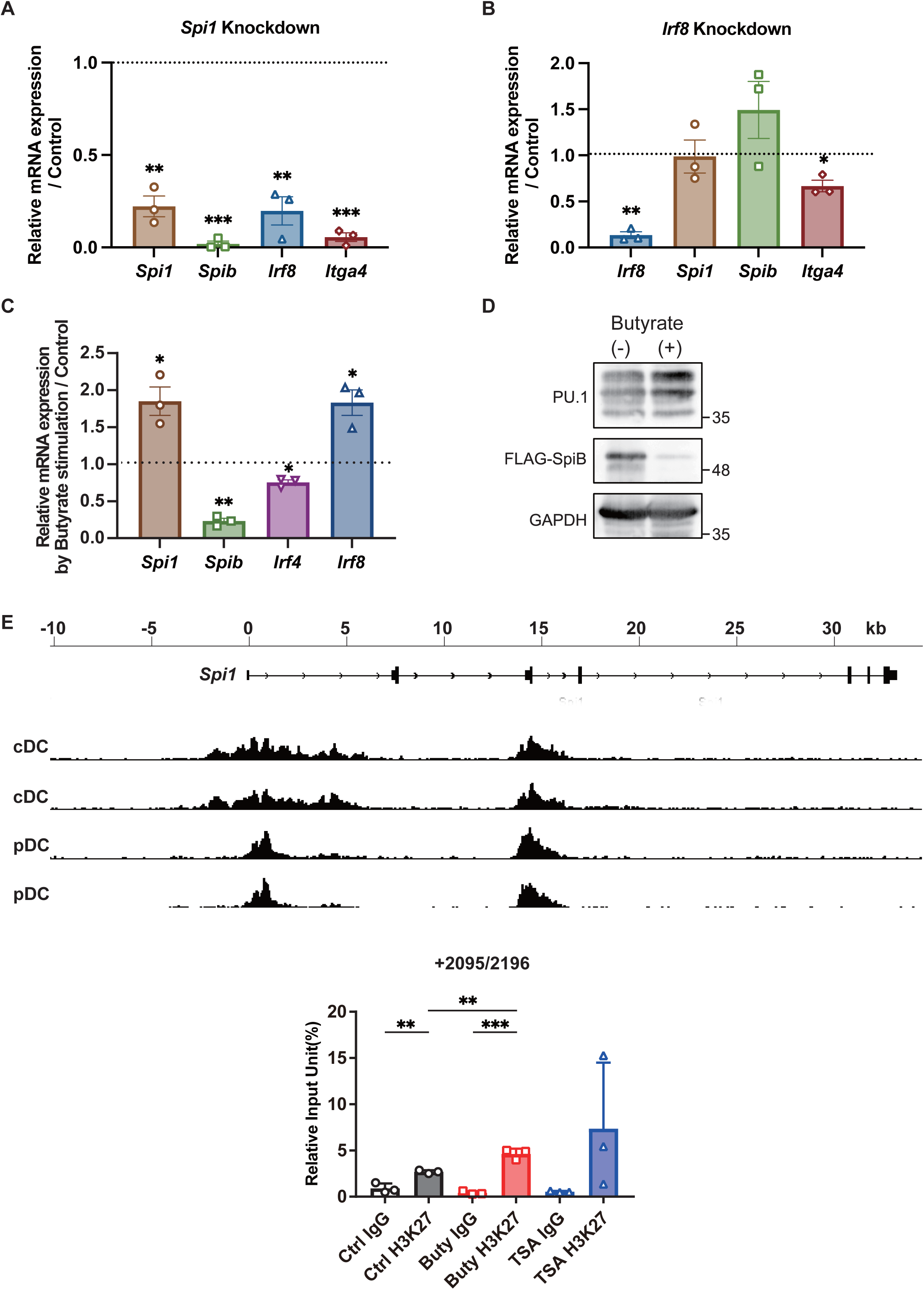
Roles of DC-related transcription factors in *Itga4* expression and effects of butyrate on the expression of DC-related transcription factors. **A.** Effects of *Spi1* (encoding PU.1) knockdown (KD) on the mRNA levels of DC-related transcription factors and *Itga4*. Introduction of siRNA into GM-CSF-induced BMDCs was performed on day 6, and after additional 48 h cultivation, BMDCs were harvested to measure mRNA levels (**A**, **B**). **B.** Effects of *Irf8* KD on mRNA levels of DC-related transcription factors and *Itga4*. **C.** Messenger RNA levels of DC-related transcription factors in butyrate-treated BMDCs. Flt3L-induced BMDCs incubated in the presence or absence of butyrate were harvested on day 8 (**C**, **D**). **D.** Western blotting profiles showing protein levels of PU.1 and SpiB in butyrate-treated BMDCs. The BMDCs were derived from 3xFlag*-Spib* mice. Whole cell lysate containing 10 μg protein was loaded to each lane. Similar results were obtained in another independent experiment. **E.** H3K27 acetylation analyzed by a ChIP assay performed with an anti-acetyl Histone H3 (Lys27) Ab and its control IgG. Top: The profiles of H3K27 acetylation around the *Spi1* gene in DCs (accession numbers are SRX1255003, 1255004, 1255005, and 1255006) were obtained via a ChIP-Atlas analysis (http://chip-atlas.org). Bottom: Relative input unit (%) of immunoprecipitated DNA assessed by qPCR targeting the indicated location on the *Spi1* gene. Relative mRNA levels, expressed as fold change relative to control (ΔΔCt method), are shown (**A**-**C**). Data represent the mean ± SEM of three independent experiments performed using biologically independent samples (**A**-**D**). The one sample *t*-test (**A**-**C**) and Tukey’s multiple comparison test (**E**) was used for statistical analyses. GM-CSF-induced BMDCs (**A**, **B**) and Flt3L-induced BMDCs (**C**, **D**, **E**) were used. *, *p* < 0.05; **, *p* < 0.01; ***, *p* < 0.005.

PU.1 is essential for the generation of DC common progenitors, and sustained PU.1 expression promotes their differentiation into cDCs, whereas reduced PU.1 expression in DC common progenitors has been shown to induce their differentiation into pDCs ^23,24^. SpiB, the amino acid sequence of which in the Ets domain exhibits high homology to that of PU.1, is involved in the development of pDCs ^25, 26^. IRF8 and IRF4 are required for the development and gene expression of cDC1 and cDC2, respectively ^27, 28^. Quantitative PCR showed that the butyrate treatment significantly increased the mRNA levels of *Spi1* and *Irf8*, and reduced the mRNA levels of *Spib* and *Irf4* in Flt3L-BMDCs (**Fig. 5C**). We confirmed that the increase in *Spi1* mRNA and decrease in *Spib* mRNA were reflected at protein level by a western blotting analysis of BMDCs derived from Flag-tagged-*Spib* knock-in-mice, which we have generated to analyze the SpiB protein in this study (**Fig. 5D**). A ChIP assay revealed that histone acetylation in the *Spi1* gene was significantly enhanced by the butyrate treatment and tended to be increased by the TSA treatment (**Fig. 5E**). Then, to elucidate the role of PU.1 in gene expression in Flt3L-induced BMDCs, we conducted a KD experiment by transfecting *Spi1* siRNA. Flow cytometric analysis revealed that *Spi1* KD significantly reduced cDC/pDC ratio (**Fig. 6A**). Although *Itga4* mRNA levels showed a tendency to decrease with *Spi1* KD, the extent of the decrease was more gradual compared to **Fig, 5A** probably due to limited *Spi1* KD degree (approximately 50 %) under these experimental conditions (**Fig. 6B**). Furthermore, we found that *Spi1* KD decreased the mRNA levels of *Irf4* and *Irf8* in Flt3L-BMDCs (**Fig. 6B**), as observed in GM-CSF-BMDCs (Ref. ^29^ and **Fig. 5A**). PU.1 positively regulated *Spib* expression in GM-CSF-BMDCs (**Fig. 5A**); however, the situation may be more complex in Flt3L-BMDCs treated with *Spi1* siRNA, as the *Spib* mRNA levels reflect an increase in the frequency of pDC and a decrease in *Spib* mRNA in cDC (**Fig. 6B**).

**FIGURE 6.**
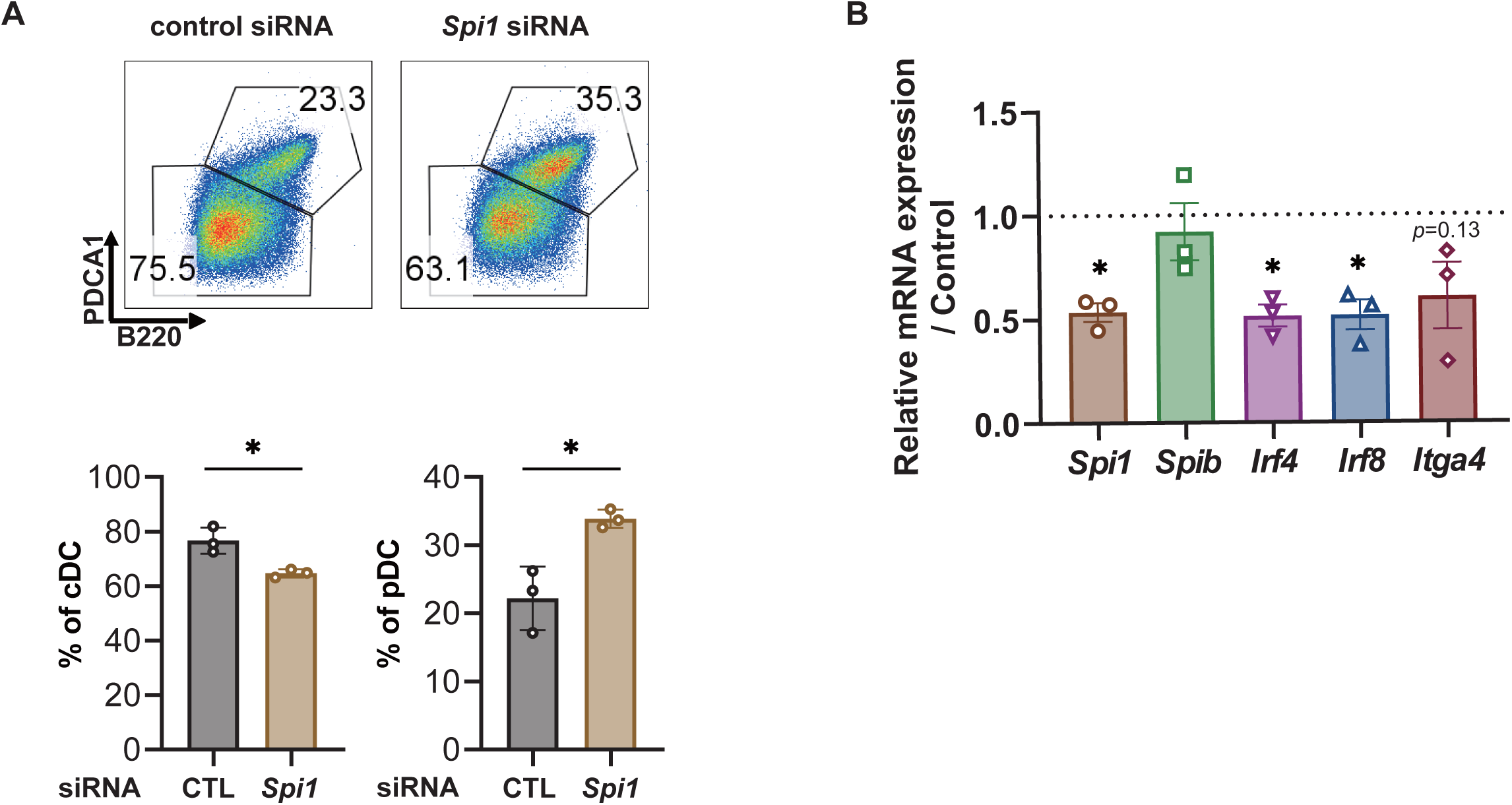
Effects of *Spi1* knock down (KD) on Flt3L-induced BMDCs. **A.** and **B.** Effects of *Spi1* KD on the expression levels of cell surface molecules (**A**) and mRNAs (**B**) in Flt3L-induced BMDCs. BMDCs transfected with *Spi1* siRNA or its control on day 6, were analyzed by a flow cytometry (**A**) and were harvested for qPCR analysis (**B**) on day 8. Data represent the mean ± SEM of three independent experiments performed using biologically independent samples (**A**, **B**). The two-tailed paired Student’s *t*-test (**A**) and the one sample *t*-test (**B**) were used for statistical analyses. *, *p* < 0.05.

Taken together, *Itga4* gene expression was dependent on PU.1 and IRF8 and butyrate transactivated the *Itga4* gene by increasing the histone acetylation of this gene in DCs. In addition, butyrate up-regulated histone acetylation in the *Spi1* gene, which is involved in the increase in the cDC/pDC ratio (**Fig. 7**).

**FIGURE 7.**
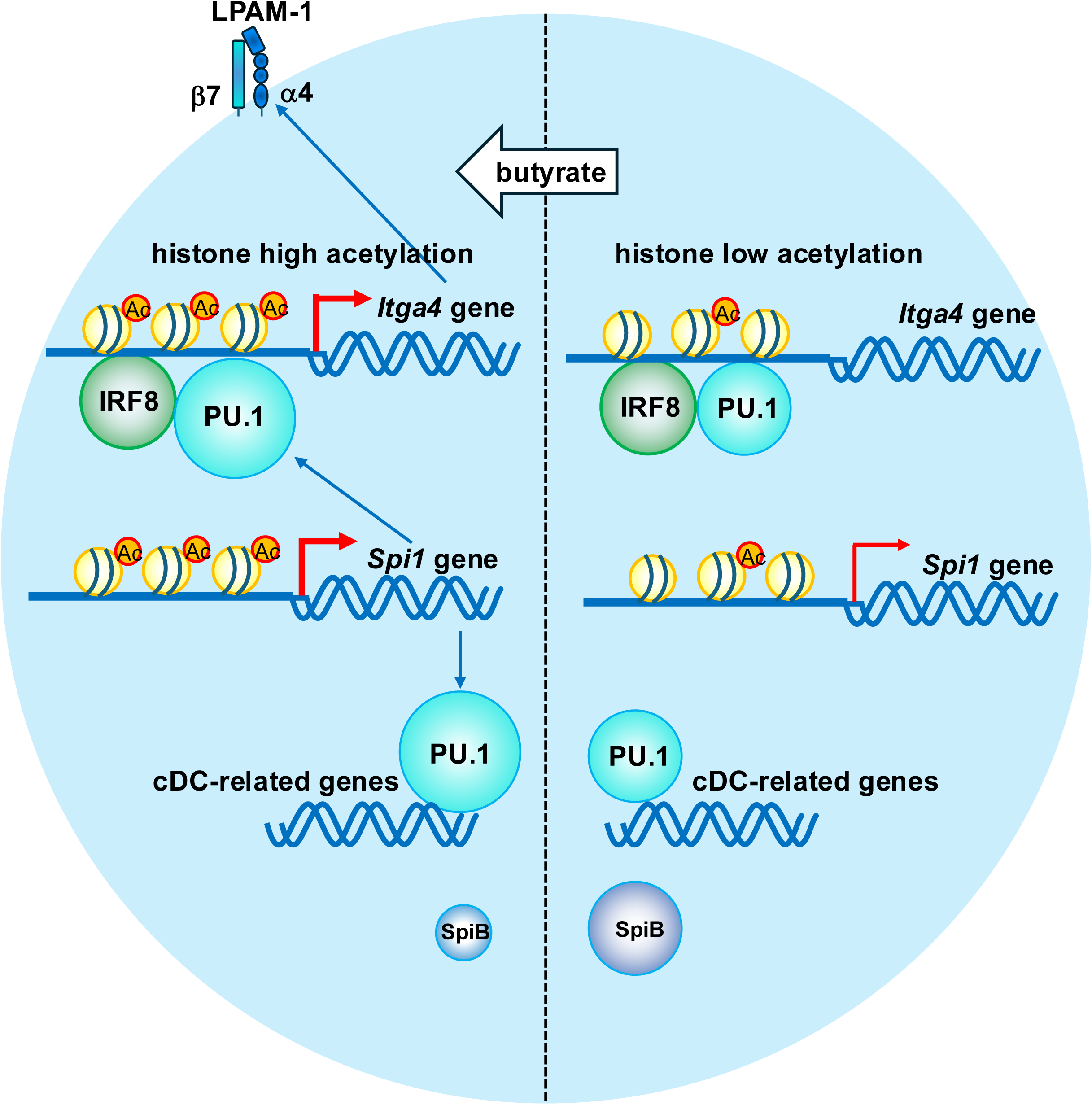
A schematic diagram illustrating the results of this study. In DCs, butyrate promotes the transcriptions of the *Spi1* and *Itga4* genes by enhancing histone acetylation of these genes through its HDAC-inhibitory activity. PU.1 and IRF8 activate the transcription of the *Itga4* gene.

## Discussion

SCFAs exert anti-inflammatory effects by modulating the functions of immuno-related cells via Gi-type GPCRs and HDACi. The present study showed that SCFAs, particularly butyrate, regulated the expression of various cell surface molecules on DCs. Briefly, SCFA treatments increased the surface levels of the antigen presentation-related molecules, MHCII and CD86, increased the cDC/pDC ratio, and induced the cell surface expression of the integrin LPAM-1 on DCs. Further detailed analyses revealed that butyrate increased the mRNA level of *Itga4* encoding the α4 subunit of LPAM-1 by enhancing the histone acetylation of the *Itga4* gene promoter with its HDACi activity, and also that the butyrate-induced up-regulation of LPAM-1 was detected in DCs in the spleen.

The involvement of HDAC in DC development was recently investigated using the cell type-specific deletion of HDAC1 or HDAC2 in mice ^30^. The present study revealed that the hematopoietic cell-specific deletion of HDAC1, but not HDAC2, impaired the development of pDC and cDC2, while not affecting cDC1 differentiation. In HDAC1-deleted cDC2, the IRF4/IRF8 ratio at the mRNA and protein levels was reduced and *Spib* mRNA levels and the H3K27 acetylation of the *Spib* gene both increased ^30^. These findings on HDAC1-deleted DCs, i.e., a reduced number of pDCs, decreases in *Irf4* mRNA, and increases in *Irf8* mRNA, are consistent with the present results on butyrate-treated DCs, suggesting that HDAC1 is a target of the HDACi activity of butyrate. The decreased mRNA level of *Spib* in butyrate-treated DCs may reflect the reduced number of pDCs. Although previous studies suggested that pDC development is dependent on HDAC, the mechanisms by which HDAC induces the development of pDCs remain unclear. Based on the present results showing *Spi1* mRNA level and histone acetylation in the *Spi1* gene were increased in butyrate-treated DCs, whereas *Spi1* siRNA transfection decreased cDC/pDC ratio, we suggested that enforced expression of the *Spi1* gene by HDACi activity of butyrate accelerated the development towards to cDCs and suppressed the commitment to pDCs. It is reported that high expression of PU.1 in DC progenitors accelerates cDC differentiation, while simultaneously suppressing pDC differentiation caused by reduced expression of PU.1^23,24^. Further studies are warranted to clarify the mechanisms by which high level of PU.1 suppresses the development of pDCs. Since we cannot rule out the possibility that the effects of butyrate on cDC/pDC ratio are due to the suppression of pDC survival and/or proliferation, even though we confirmed that 0.5 mM butyrate did not suppressed survival of splenic pDCs (data not shown), it is also necessary to examine the effects of butyrate on the survival and expansion of cDC/pDC over time.

In addition to HDAC1, HDAC3 has been suggested as a target of the HDACi activity of butyrate in DCs ^31^. This study used an originally established culture model of BMDCs from BM-derived pre-cDCs and showed that butyrate or a HDAC3 inhibitor suppressed the development of cDC2 with a decrease in the expression of IRF4 and an increase in the expression of IRF8. The decrease in IRF4/IRF8 in butyrate- or HDAC3 inhibitor-treated cDC2 was consistent with previous findings on HDAC1-deleted DCs ^30^ and the present results. However, since we only showed the mRNA expression levels of *Irf4* and *Irf8* in whole BMDCs, a more detailed analysis is required to elucidate the effects of butyrate on DC subset. This includes examining the protein expression levels of IRF4 and IRF8 in DCs while distinguishing between cDC1, cDC2, and pDC. Furthermore, although it has been reported that oral administration of butyrate affects the localization of cDCs in the small intestine in steady-state C57/BL6 mice ^31^, the involvement of integrins in this observation has not been investigated. We aim to establish experimental conditions in which SCFA administration affects DC localization in vivo, and to conduct experiments to elucidate the biological significance of SCFA-mediated DC regulation in immune-related diseases.

The previous study also showed that the butyrate treatment decreased the cell surface levels of CD80 and CD86 in cDCs ^31^. In a study on human monocyte-derived DCs and macrophages, butyrate decreased the expression of CD40, CD86, and MHCII on DCs ^32^. A study suggested that the down-regulation of PU.1 expression and PU.1 binding to the *Spi1*, *Irf8*, and *Flt3* genes in HDACi-treated DC progenitors contributed to the impaired development of pDCs ^33^. Further detailed studies on the effects of butyrate and HDACi on DCs at various developmental stages are needed to explain the reason why the discrepancy in the cell surface levels of MHCII and co-stimulatory molecules and in the role of PU.1 in pDC development between these studies and the present results.

LPAM-1 is involved in trafficking of leukocytes, including cDCs and T cells, towards the intestinal mucosa by interacting with the adhesion molecules, MAdCAM-1 and ICAM-1. The interaction between these adhesion molecules, which causes the inappropriate retention of lymphocytes in inflammatory lesions, is a therapeutic target for chronic IBD, Crohn’s disease and ulcerative colitis. Therefore, several neutralizing Abs have been developed for IBD; however, the regulatory mechanisms of LPAM-1 expression have yet to be clarified. We herein demonstrated epigenetic regulation by histone acetylation and transactivation by PU.1 and IRF8 of the *Itga4* gene, resulting in the increase in the LPAM-1 protein on the surface of DCs. Although butyrate treatment did not increase *Itgb7* mRNA levels, the observation that *Itgb7* mRNA levels in DCs were approximately 8 times higher than *Itga4* mRNA levels suggests that α4 protein levels may be the rate-limiting step in the cell surface expression of LPAM-1. Further studies are needed to confirm whether HDACi and PU.1 are commonly involved in the expression of LPAM-1 on other lymphocytes, and also to evaluate the utility of inhibitors of histone acetyltransferase and PU.1 in leukocyte trafficking, which may lead to the development of treatments for IBD. In addition, siRNAs for *SPI1/Spi1*, which suppress the chemokine-mediated migration of DCs^29^ and T cells^34^, represent other candidates. While this study focused on LPAM-1 on DCs, given that SCFAs exert potential influence on gene expression in epithelial cells ^5, 35, 36^, a further study should be conducted to clarify whether SCFAs are involved in the localization of hematopoietic cells by regulating the expression of MAdCAM-1 and ICAM-1 on epithelial cells.

It should be noted that a limitation of this study is that these results, obtained by using mouse DCs, needed to be replicated in human DCs. Clinical studies have shown that SCFAs play a crucial role in the development of the immune system shaping susceptibility to allergic disorders ^37, 38^. Furthermore, it has been established that histone acetylation has a pivotal importance in allergic diseases ^39, 40^. Therefore, it is conceivable that SCFAs exert similar effects on human DCs. We plan to conduct further experiments using human DCs to determine whether the effects of SCFAs on mouse DCs are similarly observed in humans.

## Abbreviations used

BMDC: bone marrow-derived DC;
cDC: conventional DC;
GPCR: G protein-coupled receptor;
HDAC: histone deacetylase;
KD: knockdown;
SCFA: short-chain fatty acid;
pDC: plasmacytoid DC;
PTX: pertussis toxin;
siRNA: small interfering RNA;
TSA: trichostatin A.

## Acknowledgments

This work was supported by a Grants-in-Aid for Scientific Research (B) 23K26860 (CN), 23H02167 (CN), and 20H02939 (CN); a Grant-in-Aid for Early-Career Scientists 24K17872 (KN); a Research Fellowship for Young Scientists DC2 and a Grant-in-Aid for JSPS Fellows 21J12113 (KN); a Scholarship for Doctoral Students in Immunology (from the Japanese Society for Immunology to NI); a Tokyo University of Science Grant for President’s Research Promotion (CN); the Tojuro Iijima Foundation for Food Science and Technology (CN); a Research Grant from the Mishima Kaiun Memorial Foundation (CN); a Joint Research Grant from the Research Center for Radiation Disaster Medical Science (CN); a Research Grant from the Takeda Science Foundation (CN); and the Foundation for Dietary Scientific Research (KN).

We are grateful to the members of the Laboratory of Molecular Biology and Immunology, Department of Biological Science and Technology, Tokyo University of Science, for their constructive discussions and technical support. We greatly appreciate considerations from Kimihiko Yasuda, Masako Yasuda, and the late Yayoi Yasuda.

