## Supplementary Figure S1, 2, 3, and 4 for "Short-chain fatty acids modulate the development and the cell surface molecule expression of DCs by epigenetic regulation"

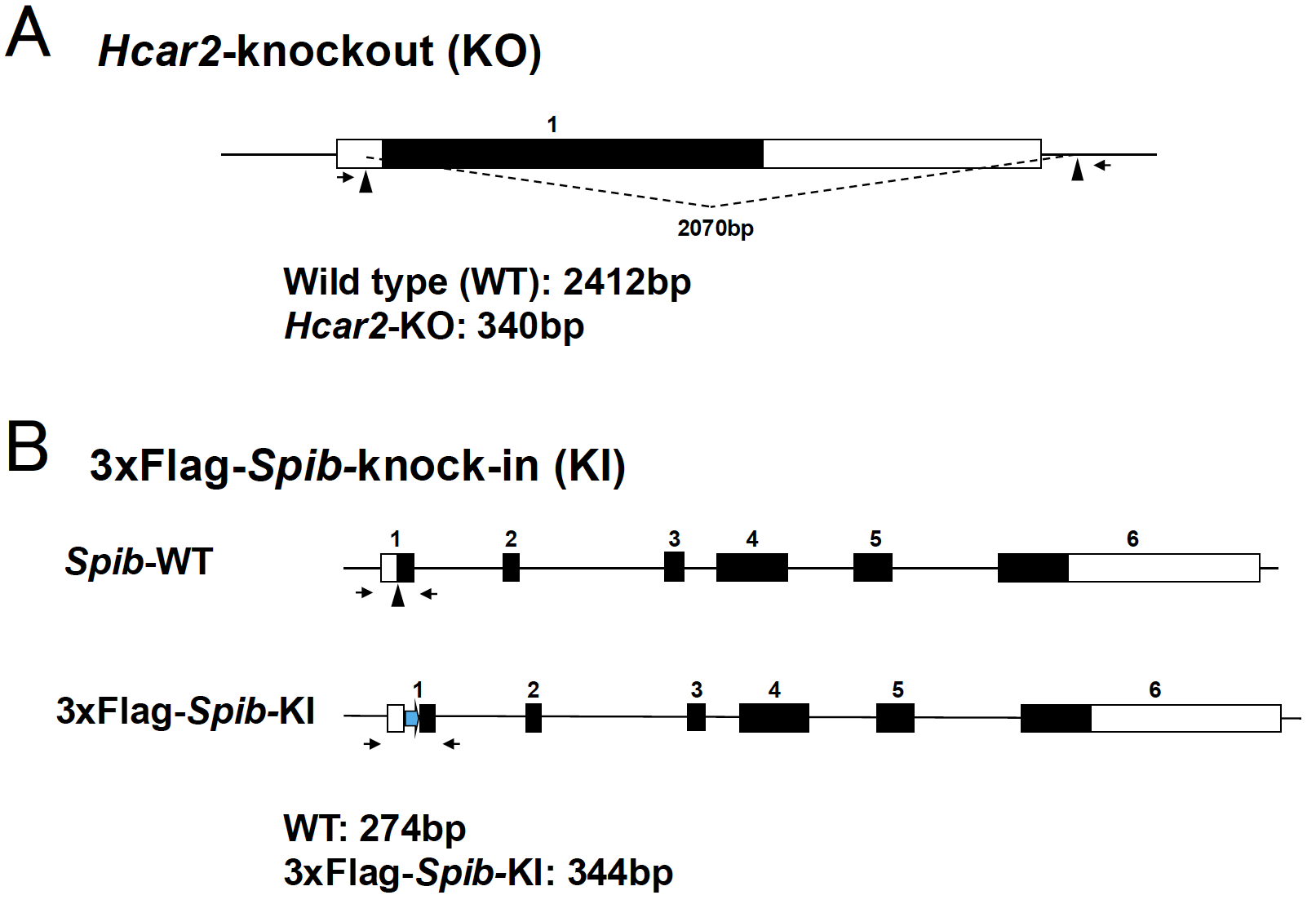

**Supplementary FIGURE S1.** Structures of the *Hcar2* gene (**A**) and the *Spib* gene (**B**) in mutant mice. Primers used for genotyping are shown as arrows.

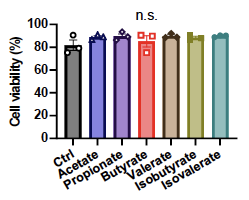

**Supplementary FIGURE S2.** Effects of short-chain fatty acids (SCFAs) on viability of bone marrow-derived dendritic cells (BMDCs).

Flt3L-induced BMDCs maintained in the presence or absence (Ctrl) of indicated SCFAs at a final concentration of 0.5 mM (from days 4 to 8) were analyzed by a flow cytometry on day 8. Cell viability was determined by DAPI staining. n.s., not significant.

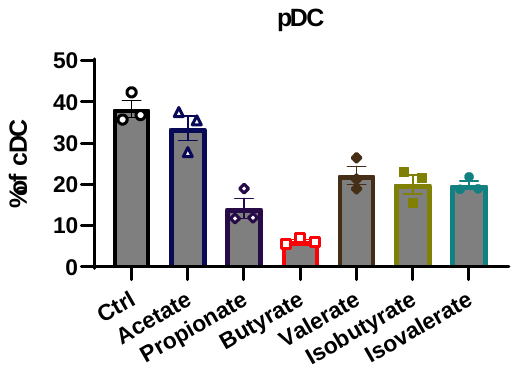

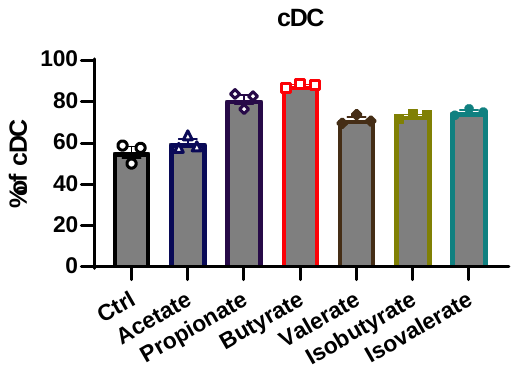

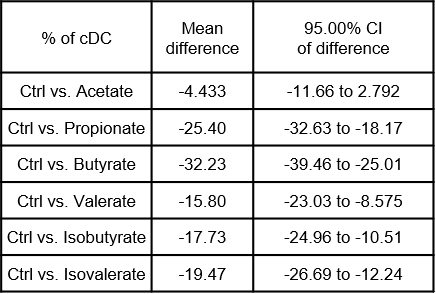

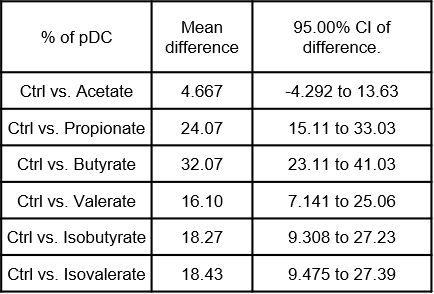

a

ab

bc

bc

c

d

d

a

a

b

b

bc

b

c

**Supplementary FIGURE S3.** Comparison of effects of short-chain fatty acids (SCFAs) on an increase in conventional DCs (cDCs; top) and a decrease in plasmacytoid DCs (pDCs; bottom).

Statistical results of **Fig. 1F** obtained by multiple comparison test were shown with letters, in which letters not shared are significantly different (left). Mean differences and their 95% confidence intervals (CIs) were also shown (right).

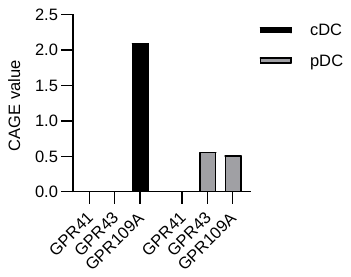

**Supplementary FIGURE S4.** Expression levels of mRNAs for GPCRs in human DCs. Data were obtained from “processed expression data of all samples for CAGE human PRJDB1099 (FANTOM5)” (https://figshare.com/articles/dataset/RefEx_expression_CAGE_all_human_FANTOM5_tsv_zip/4028613).

cDC: Dendritic Cells - monocyte immature derived,

pDC: Dendritic Cells – plasmacytoid.
